# Adipose tissue developmental growth constraints uncouple fat distribution from glucose metabolism in two mouse models of obesity

**DOI:** 10.1101/2020.08.07.239665

**Authors:** Zachary L. Sebo, Christopher Church, Ryan Berry, Matthew S. Rodeheffer

## Abstract

Subcutaneous obesity is associated with better metabolic health than visceral obesity. Here, we leverage two mouse models with differing quantities of visceral and subcutaneous fat to assess the role of fat distribution in glucose homeostasis. Interestingly, we found genetic ablation of inguinal subcutaneous fat does not exacerbate obesity-associated impairments in glucose metabolism. Consistent with this observation, mutant mice that preferentially accrue subcutaneous fat display a similar metabolic profile to controls with equal fat mass. Importantly, the increased subcutaneous adiposity in these mice occurs downstream of androgen receptor deficiency and is not driven by elevated adiponectin activity. Rather, it is caused by diminished adipocyte precursor seeding in nascent visceral fat and proportionally greater growth of subcutaneous fat. Thus, the pattern of obesogenic fat mass expansion can be determined early in development without impacting glucose metabolism. This suggests that different mechanisms underlying biased fat accumulation exert different effects on glucometabolic health.

## Introduction

The site of adipose tissue expansion in obesity is a major determinant of cardiometabolic disease risk, with high visceral adiposity associated with incident diabetes, atherosclerosis and hyperlipidemia (Carr and Brunzell, 2004, Hartz et al., 1983, Yamashita et al., 1996). Excessive visceral fat accumulation is thought to cause an elevated release of free fatty acids (FFAs) into the circulation, leading to ectopic lipid deposition and insulin resistance in peripheral organs (Björntorp, 1990, Shulman, 2014). On the other hand, high subcutaneous adiposity, specifically around the hips and thighs, is associated with a reduced incidence of metabolic disease (Hamdy et al., 2006, Karpe and Pinnick, 2015, Manolopoulos et al., 2010).

The beneficial effects of increased subcutaneous fat are thought to be associated with its capacity to function as a metabolic sink for excess lipid (Hardy et al., 2012, Tchkonia et al., 2013), which would otherwise accumulate in visceral fat and ectopically to drive insulin resistance (Shulman, 2014). This is supported by the observation that thiazolidinediones (TZDs), a family of diabetes drugs that activate PPARy, promote subcutaneous adiposity (Miyazaki et al., 2002, Mori et al., 1999, Carey et al., 2002, Yki-Järvinen, 2004). The effects of TZDs are mediated partly through the upregulation of adiponectin (Joseph et al., 2002, Kubota et al., 2006). High circulating adiponectin is associated with a greater proportion of subcutaneous fat and metabolically healthy obesity in humans (Aguilar-Salinas et al., 2008, Buemann et al., 2005, Buemann et al., 2006, Ahl et al., 2015). Furthermore, the overexpression of adiponectin in leptin-deficient mice drives the massive expansion of subcutaneous fat and improves metabolic health compared to ob/ob animals, despite super-morbid obesity (Kim et al., 2007). Parsing the metabolic effects of increased subcutaneous adiposity from adiponectin activity remains an important area of investigation.

In this study we show that PPARy conditional knockout mice that lack inguinal subcutaneous fat (murine equivalent of gluteofemoral fat)(Chusyd et al., 2016) exhibit similar glucose metabolism to animals with intact inguinal fat stores, indicating this fat depot is not necessary for normal glucose regulation. Interestingly, we found that total fat mass but not fat distribution predicts glucose tolerance in these mice. Consistent with this observation, we show that the preferential accumulation of subcutaneous fat in androgen insensitive mice has no impact on glucose metabolism compared to animals with the same total fat mass. Importantly, the increased subcutaneous adiposity in these mice is not due to adiponectin but to a dramatic reduction in visceral adipocyte precursor seeding during development. As a consequence, these mice have impaired expansion of visceral fat and preferentially accumulate subcutaneous fat in diet-induced obesity. Thus, fat distribution in obesity can be determined early in development without overt effects on metabolism. These data imply that the specific mechanism by which biased fat accumulation occurs strongly influences the role of fat distribution in metabolic health.

## Results

### Inguinal subcutaneous fat is not necessary for normal glucose metabolism in obesity

A greater proportion of subcutaneous fat, specifically around the hips and thighs, is considered protective of metabolic health in obesity (Manolopoulos et al., 2010). Thus, we hypothesized that genetic ablation of inguinal subcutaneous fat in mice would eliminate the protective effects of this depot in obesity, further impairing glucose metabolism. To prevent inguinal fat development we engineered a mouse strain (Prx1-Cre:PPARy^fl/fl^) that has PPARy conditionally knocked out in the hindlimbs (Fig. 1A, B)(Sanchez-Gurmaches et al., 2015). We fed these mice and cre-controls a SD or HFD for 30 weeks starting at 6 weeks of age followed by assessments of body composition, fat distribution and glucose metabolism. A 30-week dietary regime was chosen as wildtype mice have previously been shown to develop subcutaneous obesity from extended HFD feeding (Abreu-Vieira et al., 2015). Consistently, cre-male animals develop subcutaneous obesity on HFD, whereas cre+ animals develop visceral obesity (Fig. 1C; Supp. Fig. 1E). Cre+ mice on SD also have a more pronounced visceral fat mass bias, but total fat mass, lean mass and body weight are unaffected (Fig. 1C-F; Supp. Fig. 1E). Cre-males fed a HFD have greater body weight (Fig. 1D) and fat mass (Fig. 1F) than cre+ males but have the same amount of lean mass (Fig. 1E). The reduced total fat mass in cre+ animals can be accounted for by the absence of inguinal fat and diminished mesenteric and anterior subcutaneous fat (Supp. Fig. 1B). It is unclear why mesenteric fat mass is reduced in these mice. However, lineage tracing has shown that Prx1-cre labels a subset of anterior subcutaneous adipocytes (Sanchez-Gurmaches et al., 2015) which could explain the partial reduction in size of this depot. Similar to males, female cre+ mice do not accrue inguinal fat in obesity (Supp. Fig. 1E) yet have a more modest reduction in total fat mass compared to cre-counterparts (Fig. 1F). Brown adipose tissue mass is unaffected in cre+ animals of both sexes (Supp. Fig. 1B, D). These data show that knocking out PPARy in the Prx1 expression domain results in reduced subcutaneous fat mass while the total quantity of visceral fat in cre+ and cre-mice is similar (Fig. 1B.; Supp. Fig. 1B-D).

**Figure 1.**
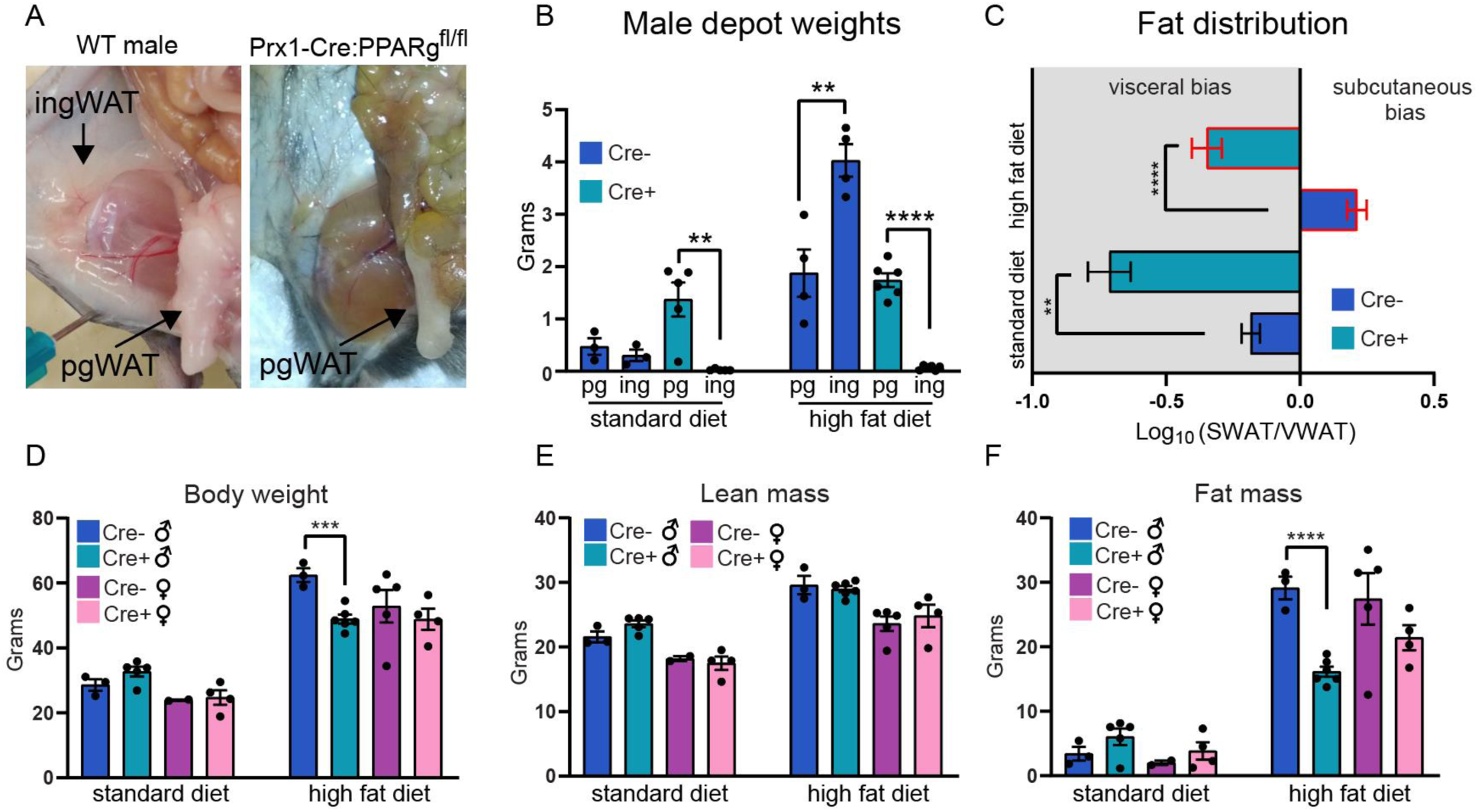
A visceral fat mass bias results from genetic ablation of inguinal subcutaneous fat. (A) Images of wildtype or Prx1-Cre:PPARy^fl/fl^ male mice at 5-6 weeks of age. (B) Perigonadal and inguinal fat depot weights in wildtype cre-males and Prx1-Cre:PPARy^fl/fl^ mice on standard diet and high fat diet (n=4-6). (C) Fat distribution of mice in (B) (n=4-6). (D) Body weight of male and female cre-/+ mice on standard diet or high fat diet (n=4-6 for males; n=2-5 for females). (E) Lean mass as determined by MRI of male and female cre-/+ mice on standard diet or high fat diet (n=4-6 for males; n=2-5 for females). (F) Fat mass as determined by MRI of male and female cre-/+ mice on standard diet or high fat diet (n=4-6 for males; n=2-5 for females). Note: Statistical significance was determined by two-tailed Student’s t-tests. Abbreviations: pg = perigonadal, ing = inguinal, SWAT = subcutaneous white adipose tissue, VWAT = visceral white adipose tissue.

Next, we assessed the impact of fat distribution on glucose metabolism in these mice. Interestingly, no differences in fasting glucose were observed between wildtype animals and those lacking inguinal fat (Fig. 2B). Neither did fasting glucose correlate with fat distribution or total fat mass (Fig. 2C, D). Indeed, age-associated hyperglycemia may override the impact of obesity (Chia et al., 2018). Surprisingly, however, the ability of Prx1-Cre:PPARy^fl/fl^ males to tolerate a glucose challenge was markedly improved relative to cre-males on HFD (Fig. 2E), despite lacking lower body subcutaneous fat. Notably, the absence of PPARy in hindlimb muscle is unlikely to impact this phenotype as glucose tolerance is unaffected in mice with PPARy knocked out in skeletal muscle (Norris et al., 2003). Given that glucose tolerance is strongly associated with total fat mass (Fig. 2G), not fat distribution (Fig. 2F), this observation can be accounted for by the reduced total fat mass in Prx1-Cre:PPARy^fl/fl^ males compared to cre-controls. Indeed, female Prx1-Cre:PPARy^fl/fl^ mice, which have a more modest reduction in total fat mass than males (Fig. 1F), do not exhibit a significant difference in glucose tolerance relative to cre-animals (Fig. 2E). Therefore, genetic ablation of inguinal fat and the resulting visceral fat mass bias do not detrimentally impact blood glucose regulation by the parameters assessed. Instead, it is the total quantity of fat, independent of its anatomic distribution, that determines glucometabolic health in this mouse model.

**Figure 2.**
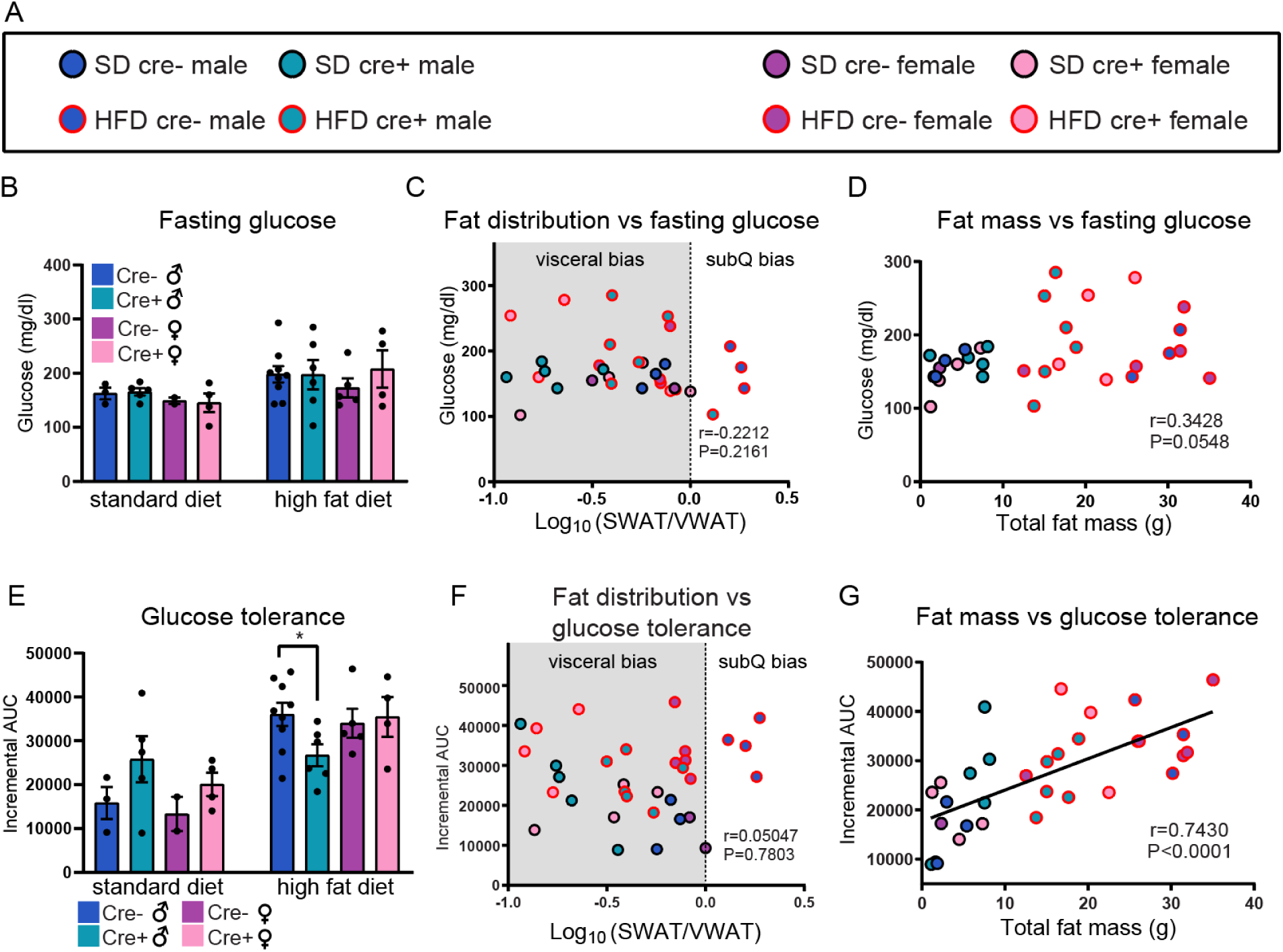
Inguinal fat ablation does not exacerbate obesity-associated impairments in glucose metabolism. (A) Legend to denote genotype and diet of animals in (C), (D), (F) and (G). (B) Fasting glucose (n= 3-9 for males; n=2-5 for females). (C) Correlation between fasting glucose and fat distribution as expressed as the logarithm of the SWAT/VWAT ratio (n=32) (D) Correlation between fasting glucose and total fat mass as determined by MRI (n=32). (E) Glucose tolerance as expressed as the incremental area under the curve (n=3-9 for males; n=2-5 for females). (F) Correlation between glucose tolerance and fat distribution (n=32). (G) Correlation between glucose tolerance and total fat mass (n=32). Note: Statistical significance was determined by two-tailed Student’s t-tests in (B) and (E). Significance in (C), (D), (F), and (G) was determined by Spearman correlation analysis. Abbreviations: SD = standard diet, HFD = high fat diet, SWAT = subcutaneous white adipose tissue, VWAT = visceral white adipose tissue.

### Increased subcutaneous fat in ARdY mice does not induce glucometabolic benefits in obesity

To further assess the impact of subcutaneous fat mass on glucometabolic health in obesity we leveraged sex-specific differences in fat distribution. Females accrue a proportionally greater quantity of subcutaneous fat and have higher adiponectin levels than males: features that are thought to contribute to the reduced risk for metabolic disease in women compared to men (Cnop et al., 2003, Geer and Shen, 2009, Palmer and Clegg, 2015). Interestingly, a subcutaneous fat mass bias is known to result from deleterious mutations in the androgen receptor (AR) gene in XY humans (Boehmer et al., 2001, Gottlieb and Trifiro, 2017, Mongan et al., 2015) and recent findings indicate similar effects occur in androgen insensitive mice (Dubois et al., 2016). Increased circulating adiponectin relative to males has also been reported for AR deficient XY (ARdY) mice (Fan et al., 2005). AR deficiency results in an anatomically intermediate phenotype between males and females such that mice and humans with this condition develop secondary sexual characteristics of females (including adiposity) yet lack internal reproductive structures other than small testes (Lyon and Hawkes, 1970, Boehmer et al., 2001, Gottlieb and Trifiro, 2017, Mongan et al., 2015). Therefore, as in previous studies with ARdY mice (Lyon and Hawkes, 1970, Yeh et al., 2002, Tanaka et al., 1994), wildtype male and female mice were used as controls for all experiments.

First, we determined the impact of diet-induced obesity on body composition and fat distribution in male, female and ARdY mice by feeding them either a SD or HFD for a period of 8 weeks starting at 7 weeks of age (Supp. Fig. 2B). At 15 weeks of age, under both dietary conditions, ARdY body weight is between that of males and females and ARdY lean mass is similar to females (Fig. 3A, B). In addition, brown fat weight in ARdY mice is no different from males or females on both diets (Supp. Fig. 4C). Notably, ARdY mice accumulate more total fat mass on SD than wildtype animals (Fig. 3C), consistent with previous findings that show an age-associated increase in fat mass in androgen insensitive mice (Sato et al., 2003). However, after 8 weeks of HFD, ARdY mice gain a similar amount of fat as wildtype animals (Fig. 3C). As expected, fat distribution in ARdY and female mice is biased toward subcutaneous depots, while male mice exhibit a visceral fat mass bias on SD (Fig. 3D). Plasma adiponectin is positively associated with subcutaneous adiposity and this is driven by the comparatively reduced adiponectin in male mice (Supp. Fig. 2C), a finding consistent with previous observations in humans (Cnop et al., 2003). Importantly, on HFD, ARdY mice preferentially accrue subcutaneous fat and wildtype animals develop a neutral fat distribution (Fig. 3D, Supp. Fig. 2D). Since high adiponectin activity has been shown to drive the preferential accumulation of subcutaneous fat in obesity (Kim et al., 2007), we compared plasma adiponectin levels in lean and obese ARdY mice. We found that plasma adiponectin is reduced in obese ARdY mice (Supp. Fig. 2E), indicating adiponectin activity does not drive the preferential deposition of subcutaneous fat in these animals.

**Figure 3.**
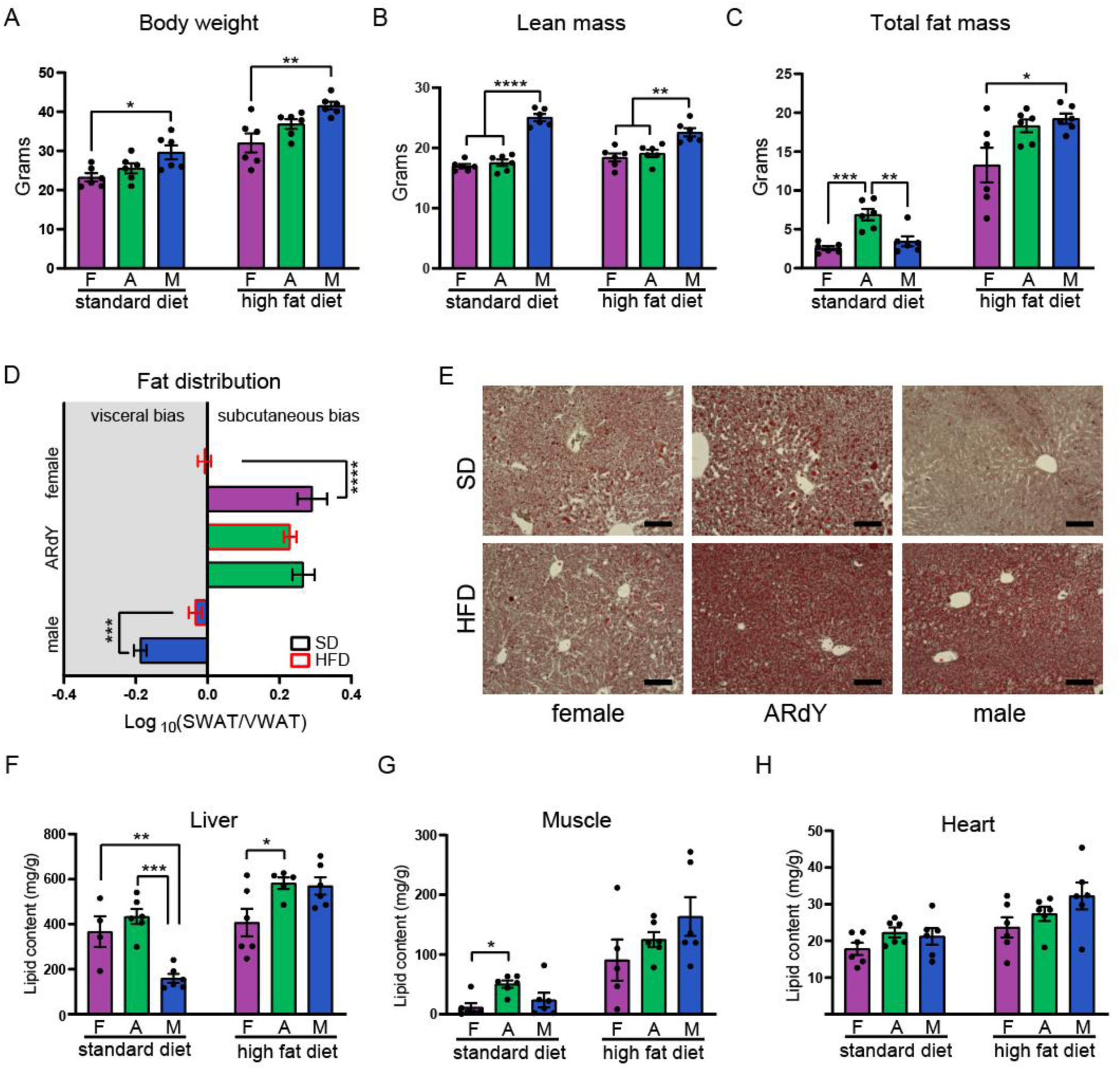
Subcutaneous obesity in ARdY mice does not protect from ectopic lipid deposition. (A) Body weight in grams (n=6). (B) Lean mass as determined by MRI (n=6). (C) Fat mass as determined by MRI (n=6). (D) Fat distribution expressed as the logarithm of subcutaneous/visceral fat mass ratio (n=6). (E) Brightfield images of oil red O stained livers from mice in (A-D). (F) Liver lipid content in mice from (A-E) (n=4-6). (G) Lipid content of gastrocnemius muscle in mice from (A-F) (n=5-6) (H) Heart lipid content in mice from (A-G) (n=6). Note: Statistical significance was determined by ordinary one-way ANOVA in (A-C) and (F-H); in (D) significance was determined by a two-tailed Student’s t-test. Scale bar in (E) is 100 um. Abbreviations: F = female, A = ARdY, M = male, SWAT = subcutaneous white adipose tissue, VWAT = visceral white adipose tissue, SD = standard diet, HFD = high fat diet

**Figure 4.**
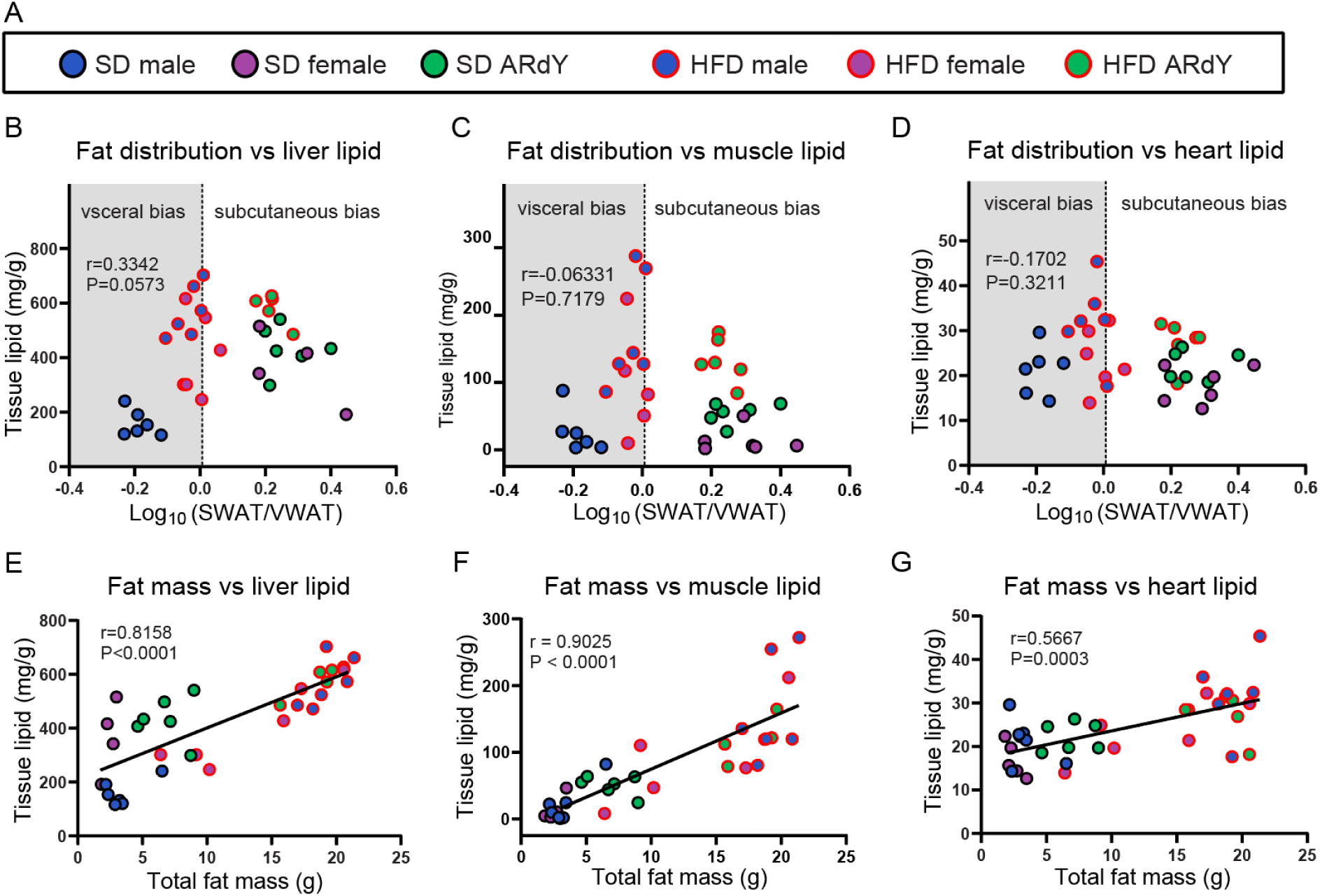
Total fat mass, not fat distribution, predicts ectopic lipid deposition in peripheral organs. (A) Legend denoting mouse genotype and diet. (B) Correlation between liver lipid content and fat distribution. (C) Correlation between gastrocnemius muscle lipid content and fat distribution. (D) Correlation between heart lipid content and fat distribution. (E) Correlation between liver lipid content and total fat mass as determined by MRI. (F) Correlation between gastrocnemius muscle lipid content and total fat mass as determined by MRI. (G) Correlation between heart lipid content and total fat mass as determined by MRI. Note: each dot represents a single animal for which tissue lipid content, total fat mass and fat distribution were determined (n=33-36 for each plot). Statistical significance was determined by Spearman’s correlation analysis.

Given that subcutaneous obesity is thought to protect from ectopic lipid deposition, we quantified the lipid content of liver, gastrocnemius muscle and heart in ARdY mice and controls. Surprisingly, ectopic lipid deposition in the livers of ARdY mice, as assessed by oil red O staining, is not improved relative to wildtype animals on SD or HFD (Fig. 3E). Consistent with the oil red O staining, a direct quantitation of lipid content in the liver, gastrocnemius and heart show no improvement in ARdY mice relative to animals with similar fat mass (Fig. 3E-H). Indeed, fat distribution does not correlate with ectopic lipid deposition in any tissues examined (Fig. 4A-C). Rather, total fat mass predicts the lipid content of liver, gastrocnemius and heart (Fig. 4D-F). Taken together, these data support the conclusion that excess lipid is not diverted from peripheral organs to subcutaneous fat in ARdY mice and show that fat distribution is not necessarily an accurate indicator of ectopic lipid deposition.

We also characterized diabetic phenotypes in these animals by performing glucose tolerance tests (GTTs). ARdY mice have impaired fasting glucose and glucose tolerance compared to wildtype animals on SD (Fig. 5A, D), probably due to increased fat mass (Fig. 3C). Consistent with this, total fat mass is an effective predictor of fasting glucose and glucose tolerance (Fig. 5D, G), as is ectopic lipid deposition (Supp. Fig. 3B-G). Fat distribution and total lean mass, on the other hand, do not correlate with either parameter of glucose metabolism (Fig. 5C, F; Supp. Fig. 3H, K). Indeed, impaired glucose metabolism is independently correlated with elevated visceral and subcutaneous fat mass (Supp. Fig. 3I, J, L, M). Thus, as fat mass increases so does ectopic lipid deposition and impaired glucose metabolism and this occurs independent of fat distribution.

**Figure 5.**
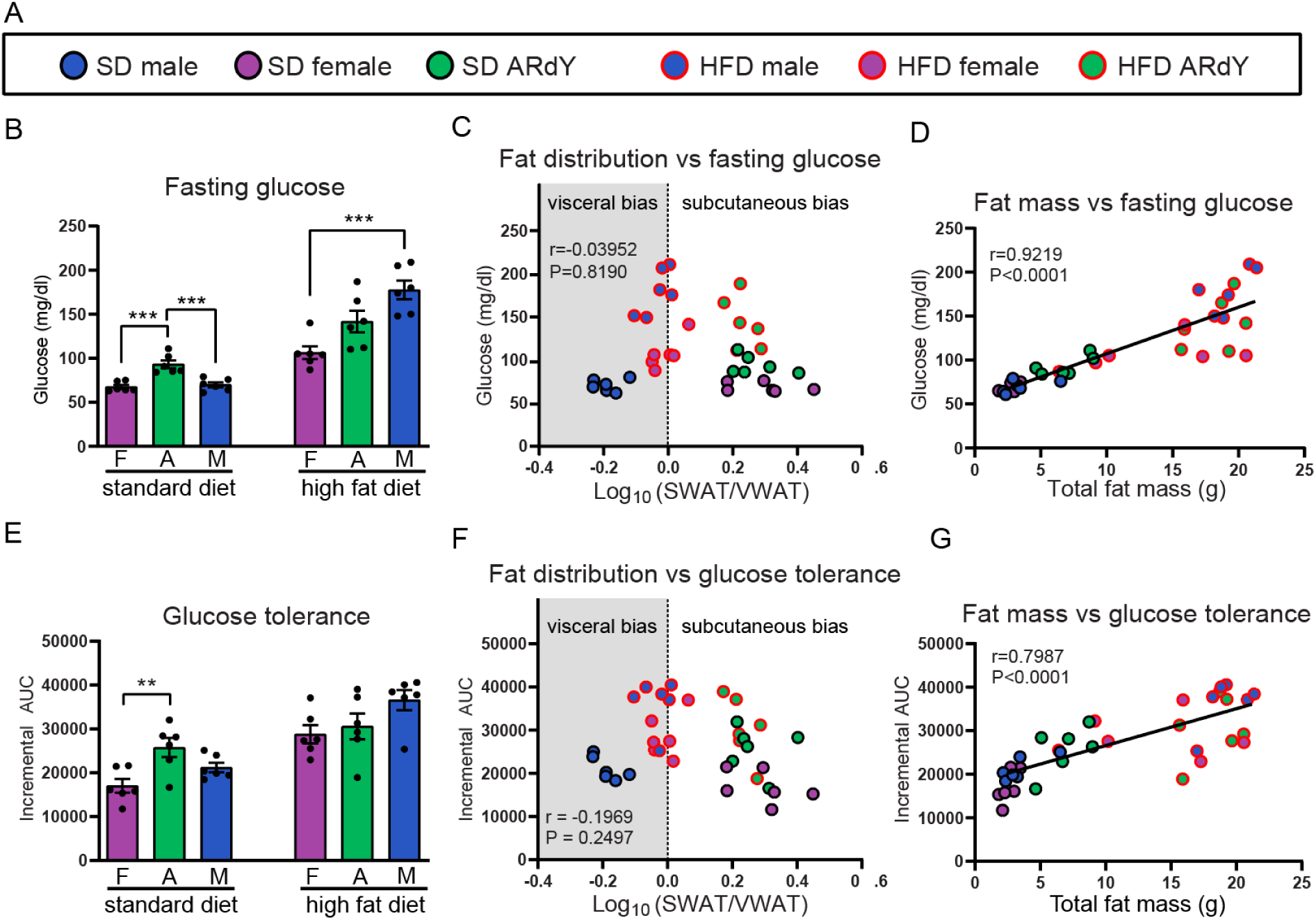
Total fat mass, not fat distribution, predicts glucometabolic health. (A) Legend denoting mouse genotype and diet. (B) Fasting glucose in 15-week-old female, ARdY and male mice on standard diet and high fat diet. (C) Correlation between fat distribution and fasting glucose. (D) Correlation between total fat mass as determined by MRI and fasting glucose. (E) Glucose tolerance expressed as incremental area under the curve. (F) Correlation between fat distribution and glucose tolerance. (G) Correlation between total fat mass as determined by MRI and glucose tolerance. Note: each dot represents a single animal for which fasting glucose and glucose tolerance were determined (n=36 for each plot). Glucose tolerance is quantitated as the incremental area under the curve. Abbreviations: F = female, A = ARdY, M = male, SWAT = subcutaneous white adipose tissue, VWAT = visceral white adipose tissue, SD = standard diet, HFD = high fat diet, AUC = area under the curve. Statistical significance in (B) and (E) was determined ordinary one-way ANOVA. Statistical significance in (C), (D), (F), and (G) was determined by Spearman correlation analysis.

Interestingly, after 8 weeks of HFD, females have significantly lower fasting glucose than males, with ARdY values directly in between (Fig. 5B). The improved fasting glucose in females is not due to reduced fat mass as lower fasting glucose is observed in female mice even when animals with less than 15g of total fat mass are excluded from analysis (Supp. Fig. 2H). Neither can the improved fasting glucose be explained by fat distribution as obese male and female mice have the same relative quantity of visceral and subcutaneous fat (Fig. 3D; Supp. Fig. 2I). Instead, fasting glucose in obese mice is inversely related to circulating adiponectin, with females and ARdY mice having greater adiponectin levels than males (Supp. Fig. 2J). These data corroborate prior studies that link improved glucose metabolism to adiponectin activity in obesity but also show that fat distribution is separable from these phenomena.

### Fat distribution is constrained by the extent of adipocyte precursor seeding in nascent adipose tissue

Given that subcutaneous obesity in ARdY mice does not improve glucose metabolism and is not caused by adiponectin activity, we sought to characterize how fat distribution is established in these animals. Interestingly, the size differential between perigonadal and inguinal fat depots is particularly pronounced in ARdY mice and is present independent of age and diet (Fig. 6A-C; Supp. Fig. 4A, B). This could be due to a depot difference in either the total number of adipocytes or the respective sizes of those adipocytes (Jo et al., 2009). We found that inguinal adipocytes are smaller than perigonadal adipocytes in ARdY mice (Fig. 6D, E), indicating the greater size of inguinal fat in these animals is due to a higher number of adipocytes in this depot.

**Figure 6.**
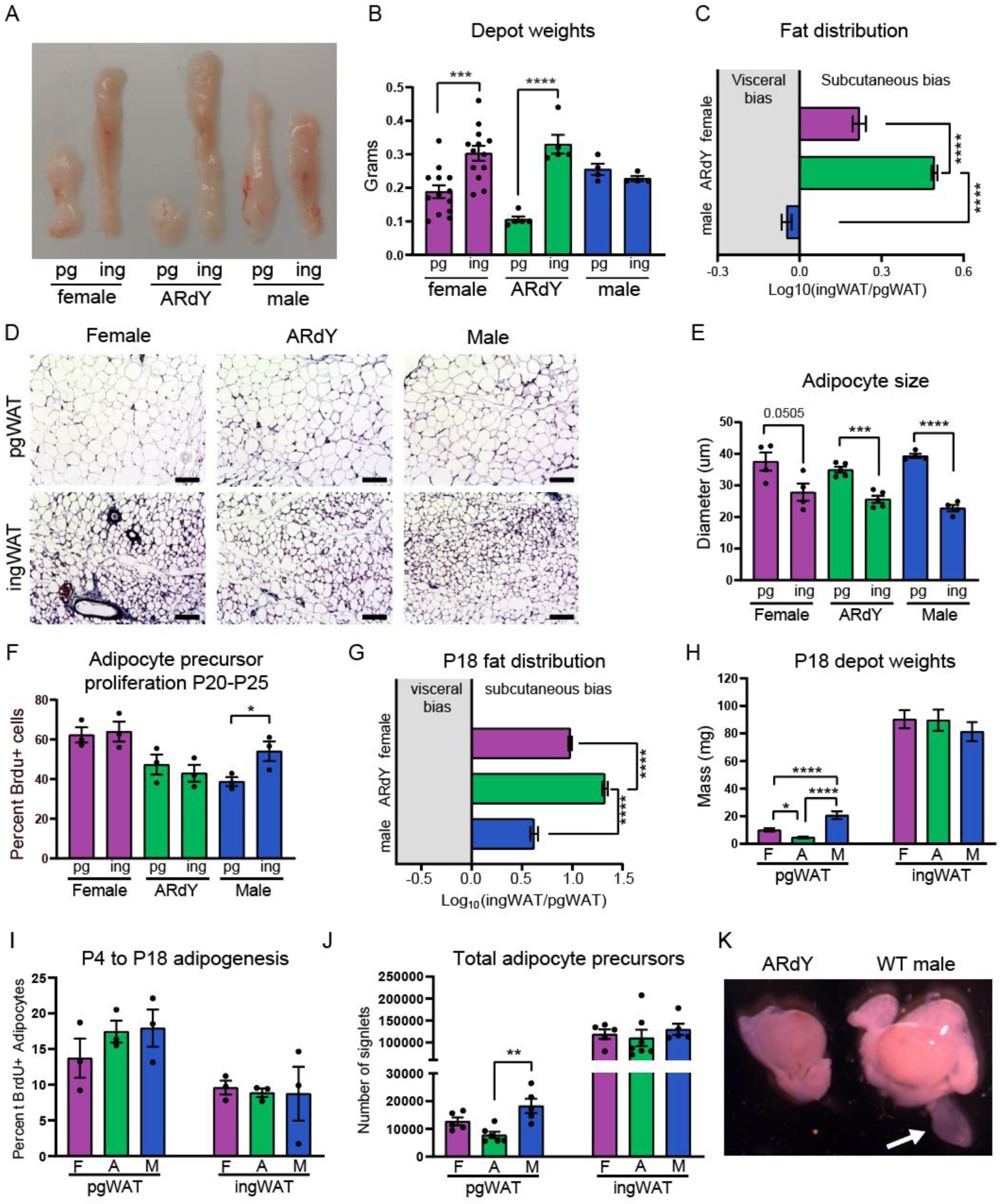
ARdY mice have a subcutaneous fat mass bias due to reduced perigonadal adipocyte precursor seeding. (A) Perigonadal and inguinal fat depots in 7-week-old female, ARdY and male mice. (B) Fat depot weights from (A)(n=4-13). (C) Fat distribution expressed as the logarithm of ingWAT/pgWAT fat mass ratio (n=4-13). (D) Trichrome stained histological sections of adipose tissue (scalebar = 100 um). (E) Adipocyte size quantification expressed as diameter (n=4-5). (F) Percentage of proliferative adipocyte precursors from P20-P25 as quantified by bromodeoxyuridine (BrdU) incorporation (cells were pooled from two animals for each genotype: technical replicates = 3, biological replicates = 6). (G) Fat distribution in P18 animals as expressed as the logarithm of the inguinal/perigonadal fat mass ratio (n=12-24). (H) Perigonadal and inguinal fat depot weights of P18 animals (n=12-24). (I) Adipocyte formation from P4-P18 as determined by BrdU pulse-chase (n-3). (J) Total number of adipocyte precursors from perigonadal and inguinal fat depots of 18-21-day old mice (n=5-7) (K) Image of testis, epididymis and epididymal appendage of ARdY and male mice at P4. Epididymal appendage is denoted by the white arrow. Note: statistical significance was determined by unpaired two-tailed Students t-tests in (B), (E) and (F). Significance in (C) and (G-J) by ordinary one-way ANOVA with Tukey’s multiple comparison’s test. Abbreviations: pg = perigonadal, ing = inguinal, SWAT = subcutaneous white adipose tissue, VWAT = visceral white adipose tissue, F = female, A = ARdY, M = male.

Total adipocyte number is determined by adipogenesis, and the rate of adipogenesis in perigonadal and inguinal fat during early puberty is thought to contribute to sex-specific fat distribution in mice (Holtrup et al., 2017). However, we found no difference in adipocyte precursor proliferation (a prerequisite to adipogenesis) between these depots from P20-P25 in ARdY mice (Fig. 6F), suggesting this developmental period does not impact fat distribution in these animals. Indeed, at P18, which is prior to the outset of puberty, ARdY mice exhibit a subcutaneous fat mass bias significantly greater than males and females (Fig. 6G). By directly comparing perigonadal and inguinal fat weights we found that the perigonadal depot is smaller in ARdY mice compared to males and females at this age, with no differences in inguinal fat weights (Fig. 6H). This indicates the subcutaneous fat mass bias in ARdY mice is due to a defect in perigonadal fat development. Therefore, we quantified adipocyte formation from E18.5 to P18 (Supp. Fig. 4H) and P4 to P18 (Fig. 6I) to identify if adipogenesis is impeded in ARdY perigonadal fat at these developmental timepoints. Surprisingly, no differences in adipocyte formation were observed between males, females and ARdY mice in either perigonadal or inguinal fat depots, indicating differential adipogenesis rates between depots does not control ARdY fat distribution.

Another factor that can impact fat mass is the total number of adipocyte precursors available to differentiate. For example, it is possible that ARdY mice have a reduced perigonadal adipocyte precursor pool such that, even with a normal rate of adipogenesis, fewer adipocytes ultimately form. Consistent with this hypothesis, ARdY mice between 18 and 21 days of age have fewer perigonadal adipocyte precursors (Fig. 6J) and total stromal vascular cells (Supp. Fig. 4F, G) than males of the same age. Moreover, it has been shown that male perigonadal fat arises from a transient structure called the “epididymal appendage” which serves as a progenitor field for the nascent fat depot (Han et al., 2011). Strikingly, the epididymal appendage is completely absent in ARdY mice (Fig. 6K). Therefore, the subcutaneous fat mass bias in ARdY mice is due to a dramatic reduction in perigonadal adipocyte precursor seeding that results in impaired perigonadal fat expansion. These results indicate that the size of the adipocyte precursor pool in nascent adipose tissue is a critical determinant of adult fat depot mass.

## Discussion

Taken together, these data show that adipose tissue develomental growth constraints can significantly impact fat distribution in adulthood without having marked effects on glucose metabolism. This is surprising given that fat distribution is typically an effective predictor of metabolic health, but is not without precedent. For example, in a mouse model of partial lipodystrophy that resembles Multiple Symmetric Lipomatosis (MSL), animals preferentially accumulate fat in the upper back while losing visceral and lower body subcutaneous fat stores. Despite this, these mice are reported to be highly insulin sensitive and glucose tolerant (Sanchez-Gurmaches et al., 2012). Consistently, impaired glucose metabolism in fatless A-ZIP/F-1 mice can be rescued by transplanting adipose tissue subcutaneously (Gavrilova et al., 2000) or by reconstituting visceral fat (Rodeheffer et al., 2008). Moreover, similar to our observations with Prx1-Cre:PPARy^fl/fl^ animals, the surgical removal of inguinal fat has no detrimental effects on blood glucose regulation in lean or obese mice (Shi et al., 2007, Foster et al., 2013).

That ARdY mice do not show improvements in ectopic lipid deposition or glucose metabolism despite developing subcutaneous obesity is likely due to the failure of subcutaneous fat to function as a metabolic sink in these animals. This suggests the channeling of excess lipid and glucose to subcutaneous fat depends on factors other than adipose tissue developmental growth constraints. Adiponectin is probably one such factor given its well-documented association with subcutaneous adiposity and improved metabolism (Kim et al., 2007, Aguilar-Salinas et al., 2008, Buemann et al., 2006, Buemann et al., 2005). Understanding the thresholds at which adiponectin promotes proper glucose regulation and subctaneous fat mass, as well as the interdependence of fat mass and adiponectin production will be important areas of future work.

The role of sex hormones in biased fat accumulation and metabolic health is also a major area of research interest (Palmer and Clegg, 2015). However, little attention has been given to the effect of these molecules on fat distribution prior to puberty. The observation that ARdY mice have impaired perigonadal fat development is intriguing given the embryonic patterning of perigonadal fat is distinct in males and females (Sanchez-Gurmaches and Guertin, 2014, Sebo et al., 2018). Thus, it is possible that androgen signaling is required not only for the proper establishment of male gonads, but for the embryonic patterning of murine perigonadal fat as well. Further work will be necessary to determine if this is the case and to identify if androgen signaling influences the embyronic establishment of human adipose tissue. Regardless, the regulation of fat distribution is unlikely to be limited to sex hormones and adiponectin. Recent large-scale GWAS studies have implicated dozens of new genes as determinants of fat distribution (Shungin et al., 2015, Heid et al., 2010) and fat distribution can be impacted by diet (Goss et al., 2013, Paniagua et al., 2007), exercise and age as well (Tchernof and Després, 2013). Thus, parsing the unique effects of individual genes, dietary components and other parameters associated with fat distribution may reveal specific drivers for the metabolic effects typically linked to visceral and subcutaneous obesity.

## Materials and Methods

### Animals and assessment of fat distribution

Animal experiments were performed according to Yale University’s Institutional Animal Care and Use Committee (IACUC). ARdY mice (Lyon and Hawkes, 1970) were obtained from Jackson Laboratories (Stock # 001809) and maintained on the C57BL/6J-A^w-j^/J background. The Eda^Ta-6J^ marker mutation was bred out of the strain prior to experiments. Prx1-Cre (Stock # 005584) (Logan et al., 2002) and PPARy^fl/fl^ (Stock # 004584) (He et al., 2003) mice were bred to generate Prx1-Cre:PPARy^fl/fl^ mice that lack inguinal fat. Standard diet in this study was made by Harlan Laboratories (2018S) and high fat diet by Research Diets (D12492). Body composition of mice was determined via magnetic resonance imaging (EchoMRI-100H, EchoMRI, Houston, TX, USA).

Fat distribution was calculated using the following equation,

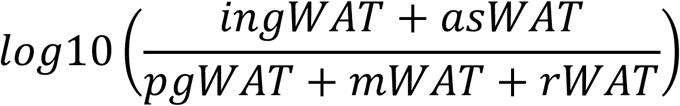

where the weight in grams of each subcutaneous (ingWAT and asWAT) and visceral (pgWAT, mWAT and rWAT) depot were combined and divided to acquire an SWAT/VWAT ratio. The logarithm of this ratio is required to correct for skew. Notably, “inguinal fat” in this study refers to the continuous adipose depot that runs along the dorsolumbar-gluteal axis of the hindlimb in mice. Only white adipose depot weights were included in the fat distribution calculation.

### Adipocyte hyperplasia and hypertrophy

Quantification of adipocyte precursor proliferation and adipocyte formation was performed as described previously (Jeffery et al., 2015, Berry and Rodeheffer, 2013). Briefly, to quantify adipocyte formation from E18.5 to P18, pregnant dams were injected intraperitoneally with 50mg/kg BrdU in PBS. Adipocyte nuclei from pups of such litters were analyzed for BrdU incorporation at P18. Similarly, to quantify adipocyte formation from P4-P18, P4 pups were injected intraperitoneally with 50mg/kg BrdU in PBS in the morning and evening and allowed to develop until P18, at which point BrdU incorporation into adipocyte nuclei was quantified. At least 100 BrdU+ adipocytes were counted for each depot. Adipocyte size was determined from images of trichrome-stained histological sections of adipose tissue using Cell Profiler (Carpenter et al., 2006). Histological sectioning and trichrome staining was performed by the Histology Core Laboratory of the Yale School of Medicine, Department of Comparative Medicine. Images were taken using a Leica SP5 confocal microscope for the quantification of BrdU+ adipocyte nuclei and a Keyence BZ-X800 microscope for adipocyte sizing.

### Glucose tolerance tests and lipid quantification

Mice were fasted overnight (16-18 h) and fasting blood glucose level was obtained via a tail vein nick. Mice were injected intraperitoneally with a 20% glucose solution in saline at 2g/glucose/kg body weight and blood glucose was measured at 10, 20, 30, 60 and 120 minutes afterward. Glucose tolerance is defined as the incremental area under the curve (i.e. the area under the curve normalized to fasting glucose). Tissue lipid quantification in liver, gastrocnemius and heart was performed using the Cell Biolabs Fluorometric Lipid Quantification Kit (Catalog # STA-617) according to manufacturer’s instructions with reagent volumes halved in technical singlicate. For Oil Red O staining, the medial lobe of the liver was fixed in 4% paraformaldehyde for ∼24 h followed by incubation in 30% sucrose in saline for ∼24 h. Livers were then embedded in OCT compound (Tissue-Tek, product code 4583) and flash frozen in liquid nitrogen. Histological sectioning and Oil Red O staining was performed by the Histology Core Laboratory of the Yale School of Medicine, Department of Comparative Medicine. Brightfield images of histological sections were taken using a Keyence BZ-X800 microscope.

### Statistical Analysis

Statistical tests were performed using GraphPad Prism (version 8.3.0). P-values < 0.05 were considered significant. Specific statistical tests used for each experiment and the number of biological replicates are denoted in figure legends. Error bars represent mean ± SEM. *P<0.05, **P<0.01, ***P<0.001, ****P<0.0001. Lines of best fit are shown only on plots with statistically significant correlations.

## Acknowledgments

The authors thank Michael Schadt and the Histology Core Laboratory for assistance with tissue sectioning and staining. We also thank Caroline Zeiss for sharing microscopy equipment and the Yale Flow Cytometry Core for equipment usage. This work was supported by a National Science Foundation Graduate Research Fellowship (DGE1122492 to Z.L.S.) and the NIDDK (R01DK110147 and DK090489 to M.S.R.).

## Supplementary Figures

**Supplementary Figure 1.**
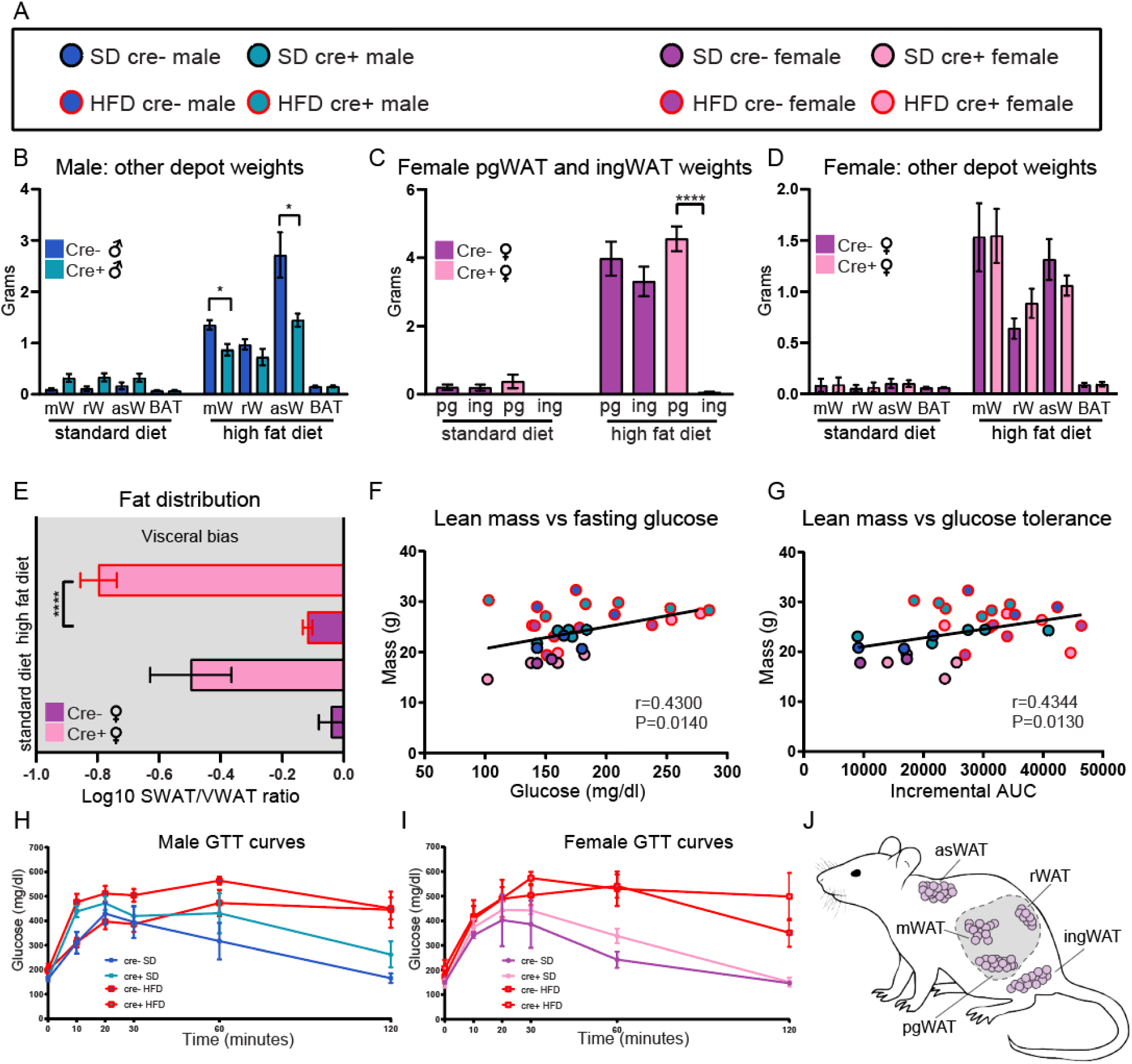
Additional fat distribution and glucose metabolism parameters in Prx1-Cre:PPARy^fl/fl^ mice. (A) Legend to denote the genotype and diet of each animal in (F) and (G). (B) Fat depot weights of male mice not shown in Figure 1B. (C) Perigonadal and inguinal fat weights in female mice (n=2-5) (D) Fat depot weights of female mice not shown in (C) (n=2-5). (E) Fat distribution of female mice (n=2-5). (F) Correlation between total lean mass and fasting glucose (n=32). (G) Correlation between total lean mass and glucose tolerance (n=32). (H) Glucose tolerance test of male mice fed HFD or SD (n=3-9). (I) Glucose tolerance test of female mice fed HFD or SD (n=2-5). (J) Schematic showing visceral and subcutaneous fat depots in mice. Note: Statistical significance was determined by two-tailed Student’s t-tests in (B), (C) and (E) and by Spearman correlation analysis in (F) and (G).

**Supplementary Figure 2.**
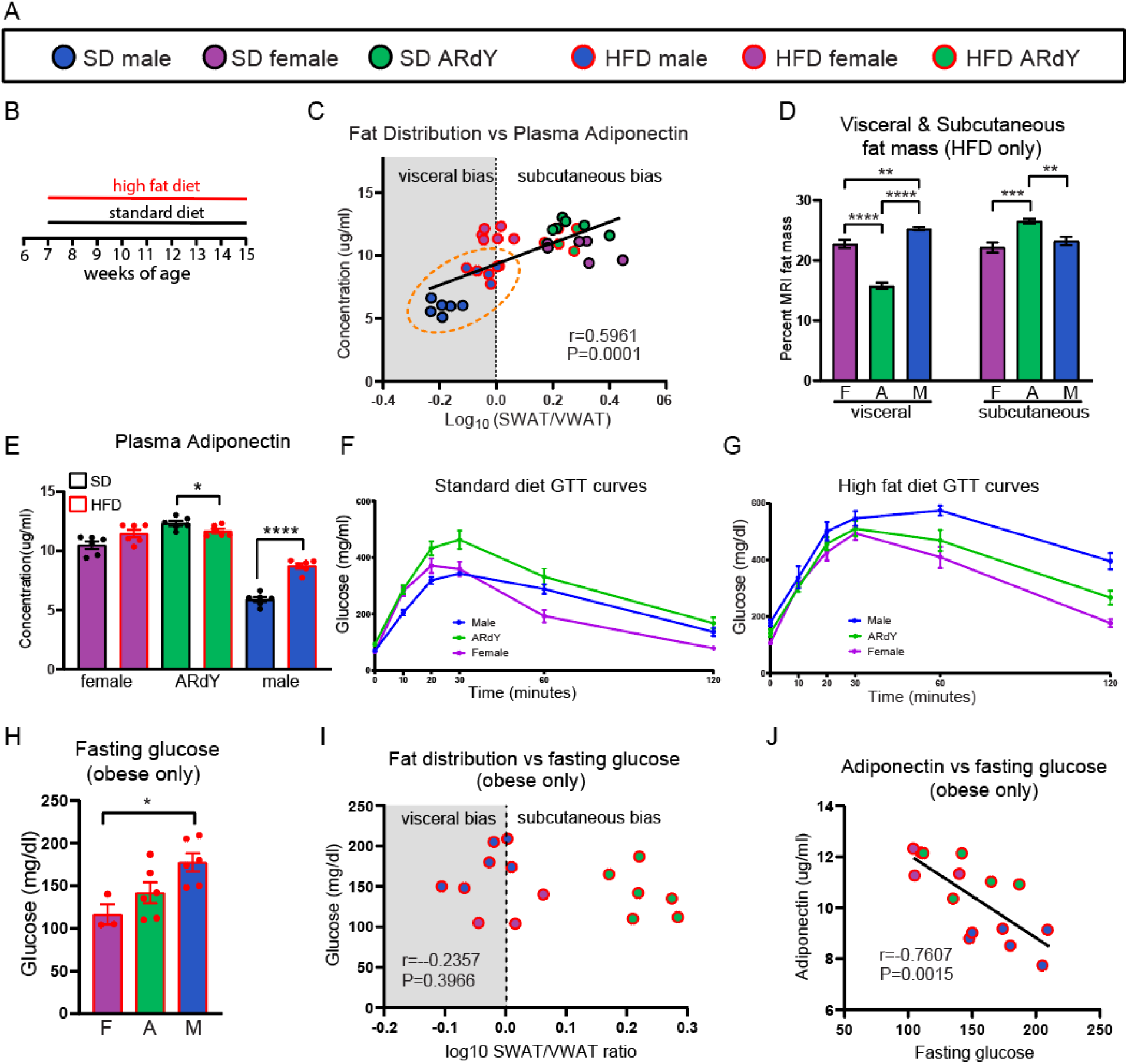
Relationships between fat distribution, adiponectin and glucose metabolism. (A) Legend to denote the genotype and diet of each animal in (C), (I) and (J). (B) Schematic showing dietary regime for male, female and ARdY mice. (C) Correlation between fat distribution and plasma adiponectin level (n=35). (D) Visceral and subcutaneous fat mass expressed as a percentage of total fat mass measured by MRI (n=6). (E) Plasma adiponectin level (n=6). (F) Glucose tolerance test of 15-week-old male, female, and ARdY mice fed standard diet (n=6). (G) Glucose tolerance test of 15-week-old male, female, and ARdY mice fed high fat diet (n=6). (H) Fasting glucose in mice with at least 15g total fat mass (n=3-6). (I) Correlation between fat distribution and fasting glucose in obese mice (n=14). (J) Correlation between plasma adiponectin and fasting glucose in obese mice (n=14). Note: Statistical significance in (B), (I) and (J) was determined using Spearman’s correlation analysis. In (D) and (H) by an ordinary one-way ANOVA with Tukey’s multiple comparison’s test and in (E) by a two-tailed Student’s t-test.

**Supplementary Figure 3.**
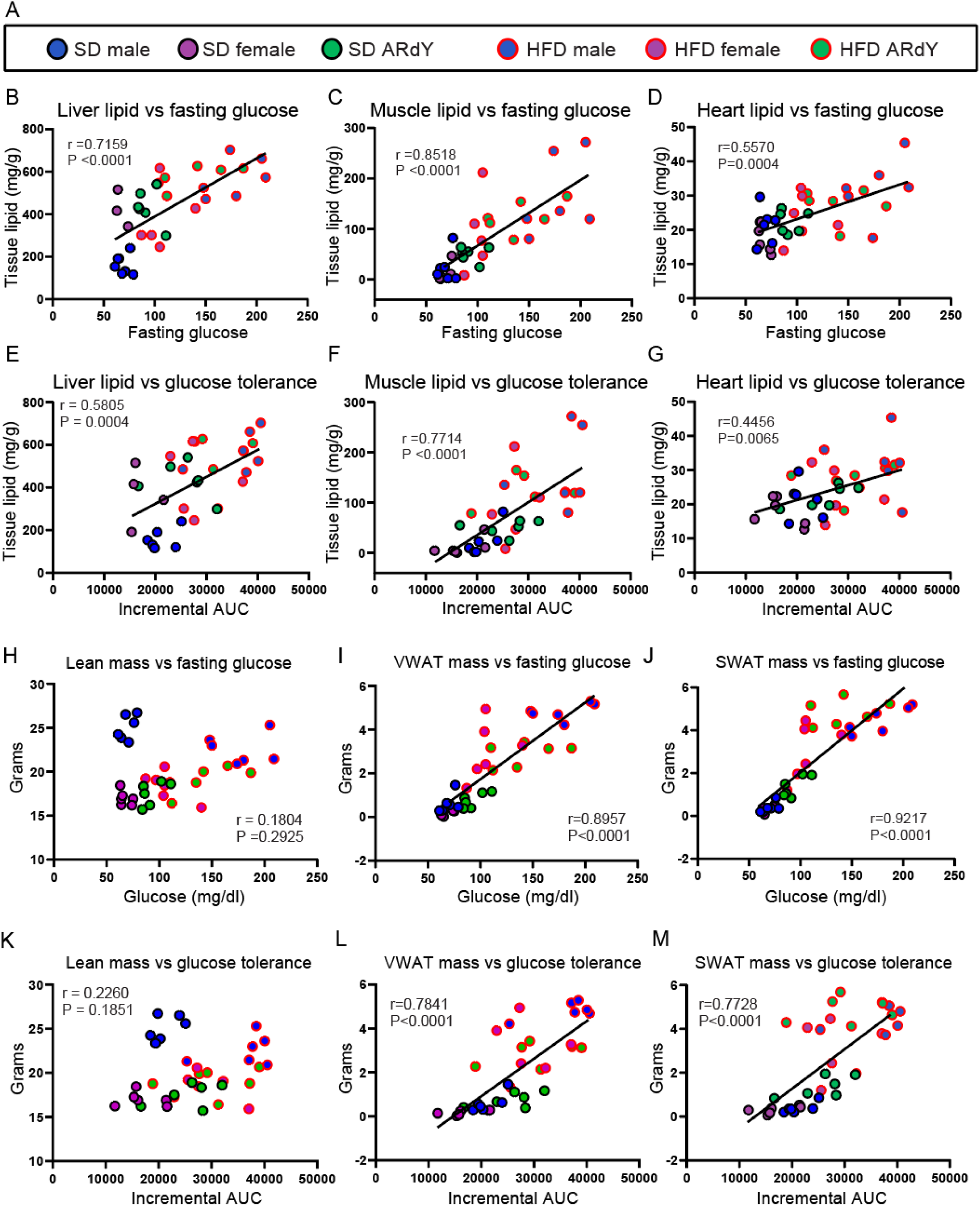
Ectopic lipid deposition and fat mass predict impaired fasting glucose and glucose tolerance. (A) Legend to denote the genotype and diet of each animal in (B-M). (B) Correlation between liver lipid content and fasting glucose (n=33). (C) Correlation between muscle lipid content and fasting glucose (n=34). (D) Correlation between heart lipid content and fasting glucose (n=36). (E) Correlation between liver lipid content and glucose tolerance (n=33). (F) Correlation between muscle lipid content and glucose tolerance (n=34). (G) Correlation between heart lipid content and glucose tolerance (n=36). (H) Correlation between total lean mass and fasting glucose (n=36). (I) Correlation between VWAT mass and fasting glucose (n=36). (J) Correlation between SWAT and fasting glucose (n=36). (K) Correlation between total lean mass and glucose tolerance (n=36). (L) Correlation between VWAT mass and glucose tolerance (n=36). (M) Correlation between SWAT mass and glucose tolerance (n=36). Note: Statistical significance was determined using Spearman correlation analysis.

**Supplementary Figure 4.**
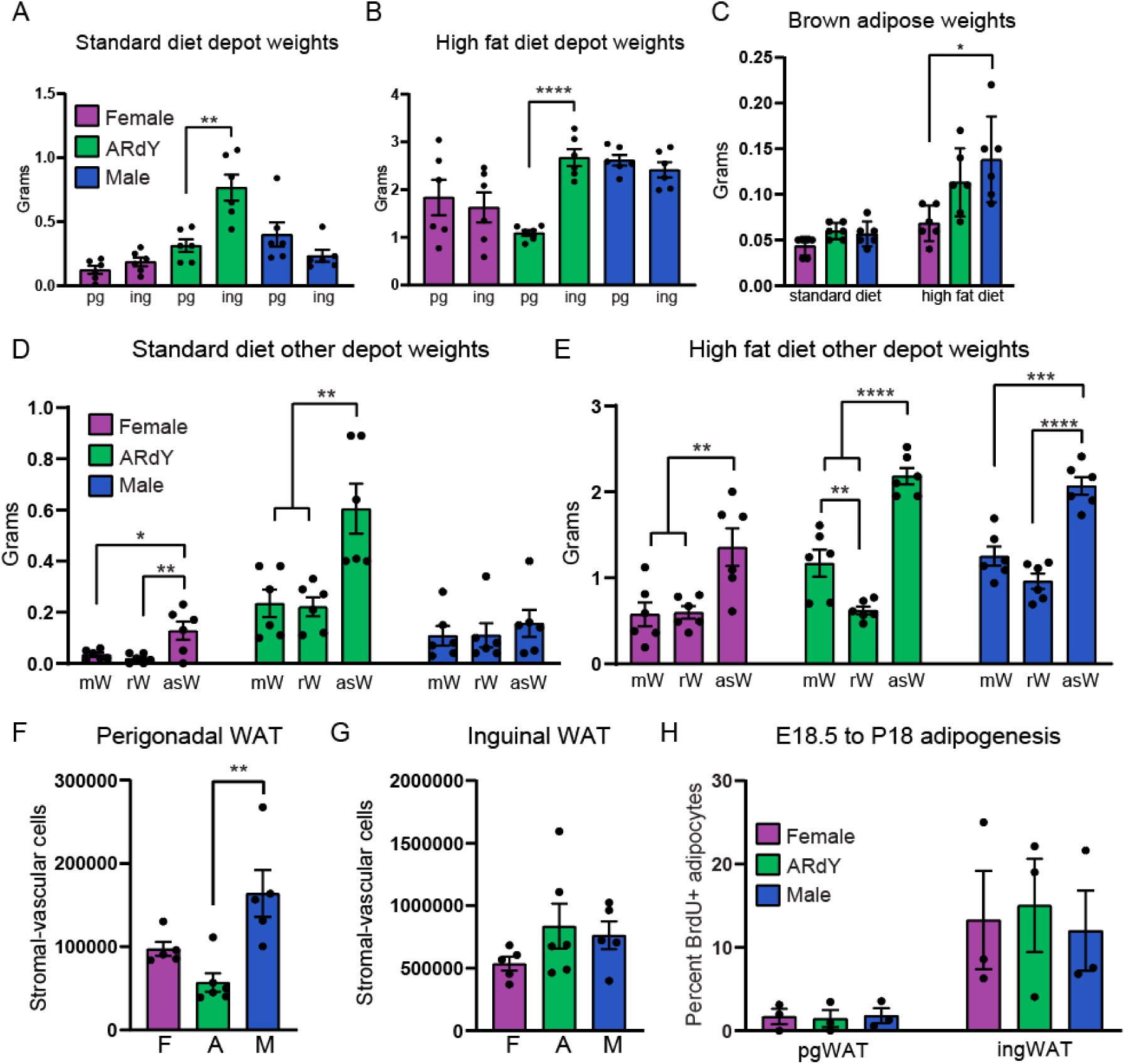
Additional depot weights and adipogenesis in female, male and ARdY mice. (A) pgWAT and ingWAT weights on standard diet (n=6) (B) pgWAT and ingWAT weights on high fat diet (n=6). (C) mWAT, rWAT, asWAT and BAT depot weights on standard diet (n=6). (D) mWAT, rWAT, asWAT and BAT depot weights on high fat diet (n=6). (E) total stromal vascular cells in pgWAT in 18-21 day old mice (n=5-6). (F) total stromal vascular cells in ingWAT in 18-21 day old mice (n=5-6). (G) New adipocyte formation from E18.5 to P18 in pgWAT and ingWAT (n=3). Note: Statistical significance was determined using unpaired Student’s t-tests in (A) and (B) and one-way ANOVA with Tukey’s multiple comparison’s test in (C-G). pg = perigonadal, ing = inguinal, mW = mesenteric, rW = retroperitoneal, asW = anterior subcutaneous, BAT = brown adipose tissue.

